# TCF1^lo^ CD8 T cells proliferate and persist autonomously in tumors

**DOI:** 10.64898/2026.01.17.700120

**Authors:** Megan M. Erwin, Natalie R. Favret, Claudia N. McDavid, Zachary D. Ewell, Melissa M. Wolf, Lincoln A. Brown, Jessica J. Roetman, Michael W. Rudloff, Kristen A. Murray, Carlos R. Detrés Román, Mary Philip

**Affiliations:** Department of Medicine, Division of Hematology and Oncology, Department of Pathology, Microbiology, and Immunology, Vanderbilt School of Medicine, Nashville, TN, USA; Vanderbilt Center for Immunobiology, Vanderbilt University Medical Center, Nashville, TN USA; Vanderbilt-Ingram Cancer Center, Vanderbilt University Medical Center, Nashville, TN, USA; Vanderbilt Institute for Infection, Immunology, and Inflammation, Vanderbilt University Medical Center, Nashville, TN USA

## Abstract

Cancers develop in humans over months to years, and tumor-specific CD8 T cells (TST) can interact with cancer cells throughout tumorigenesis. Nevertheless, the long-term population dynamics of TST, especially within progressing tumors, are not well understood. A paradigm first established in chronic viral infection and applied to tumors describes a population hierarchy among exhausted T cells. Progenitor/stem-like exhausted T cells, which express the transcription factor T cell factor 1 (TCF1), maintain the population through self-renewal and by giving rise to terminally differentiated TCF1^lo^ progeny. This has led to a focus on TCF1^hi^ T cells, and though TCF1^lo^ CD8 T cells are the predominant tumor-infiltrating/tumor-reactive subtype in patients, they have been largely overlooked. We leveraged our autochthonous liver cancer model to analyze TST differentiation and proliferation throughout tumorigenesis. Dual EdU/BrdU labeling studies revealed that throughout tumorigenesis, a subset of TCF1^lo^ TST in the liver stochastically entered and exited cell cycle, and at later time points there was no evidence of a TCF1^hi^ progenitor-like population. Moreover, TCF1-knockout TST proliferated and persisted robustly in tumors. Using liver cancer and melanoma models, we showed that tumor-resident TCF1^lo^ TST proliferate and persist autonomously, even when new TST influx into tumors is inhibited. The prevailing notion is that only TCF1^hi^ TST self-renew but we now demonstrate, using a clinically relevant mouse cancer model, that TCF1^lo^ TST stochastically proliferate to achieve long-term population maintenance. Future studies to understand and harness this mechanism to improve T cell persistence in tumors could lead to novel immunotherapies for patients with cancer.

**SYNOPSIS:** We show that tumor-specific T cells with little/no expression of TCF1, previously considered incapable of self-renewal, can proliferate stochastically and persist long-term. As TCF1^lo^ CD8 T cells are often the predominant tumor-reactive T cells found in tumors, future studies should be aimed at reprogramming these proliferating T cells within tumors.

## INTRODUCTION

Tumor-specific CD8 T cells (TST) are critical mediators of anti-tumor immunity, capable of recognizing and eliminating malignant cells. While immunotherapies harnessing T cells, including anti-PD1 blocking antibody (Immune Checkpoint Blockade; ICB) and adoptive T cell therapy with chimeric antigen receptor T (CAR T) therapies, most patients still fail to achieve durable response (1). A major barrier for successful immunotherapy is the failure of TST to persist (2,3). Thus, understanding the mechanisms underlying T cell persistence in tumors is of paramount importance.

The current paradigm for understanding T cell persistence in the setting of chronic antigen exposure came from studies in chronic viral infection. John Wherry’s group first showed that the population of virus-specific CD8 T cells follow a progenitor-progeny hierarchy of proliferation and differentiation (4). Progenitor/stem-like exhausted T cells express T cell factor 1 (TCF1; encoded by *Tcf7*), a transcription factor expressed in naive and memory T cells and associated with quiescence/stemness (reviewed in (5)). Progenitor exhausted T cells proliferate to self-renew and differentiate into more effector-like progeny, downregulating TCF1 in the process (6–8) and reviewed in (9). This paradigm has been extended to other settings of chronic antigen exposure, including tumors (10,11) and autoimmunity (12).

Progenitor TCF1^hi^ exhausted T cells were shown to preferentially proliferate in response to anti-PD1/PDL1 antibody therapy (7,8), leading to increased focus on TCF1^hi^ progenitor exhausted subsets. While some studies have shown an association between the expression of *Tcf7* in T cells and ICB responses in patients with melanoma (13–15), more recent studies in patients with renal cell carcinoma failed to find such an association (16). Most studies on TCF1-mediated progenitor/progeny hierarchies in tumors have been carried out in transplantable tumor mouse models, which are limited by short study windows between tumor implantation and endpoint (3-4 weeks). In contrast, in humans and mice, autochthonous cancers progress on the scale of months to years, and we and others have shown that the number of TCF1^hi^ progenitors declines over time during tumor progression (17), in murine and human chronic infection (18,19) and with aging (20). Moreover, in many patients, the majority of tumor-reactive CD8^+^ tumor-infiltrating lymphocytes (TIL) are TCF1^lo^ (16,21). Thus, we set out to understand the mechanisms underlying TST persistence during tumorigenesis using a clinically relevant liver cancer mouse model in which we can study TST over months. We found that later during tumorigenesis, the majority of TST in the SLO and within tumors were TCF1^lo^. Nevertheless tumor-resident TCF1^lo^ TST enter and exit cell cycle stochastically to autonomously and stably maintain the tumor-resident population long-term.

## RESULTS

### Tumor-specific CD8 T cells persist throughout tumorigenesis

To investigate longitudinal TST population dynamics in tumor-bearing hosts, we utilized our established autochthonous mouse model of inducible liver cancer (AST;Cre-ER^T2^). In AST;Cre-ER^T2^ mice, a single dose of tamoxifen causes hepatocytes to undergo Cre recombinase-mediated induction of the SV40 large T antigen (TAG), under control of the albumin promoter/enhancer (22). TAG serves as both an oncogenic driver (through inhibition of TP53 and RB) (23) and a tumor-specific antigen. We adoptively transferred naive TAG-specific CD8 T cells (TCR_TAG_) into tamoxifen-treated AST;Cre-ER^T2^ mice; in parallel, we transferred TCR_TAG_ into C57BL/6 (B6) mice infected with TAG epitope-expressing *Listeria monocytogenes* (LM_TAG_) (**Fig. 1A**). We analyzed TCR_TAG_ from the spleens and livers of infected and tumor-bearing mice at time points from 2.5 to 70 days after transfer. TCR_TAG_ were uniformly activated in the spleens and livers of infected- and tumor-bearing mice, as demonstrated by CD44 upregulation (**Supplementary Fig. 1A**) and early expansion in numbers (**Fig. 1B**). Following initial expansion, TCR_TAG_ numbers in the spleens and livers of infected mice contracted, while TCR_TAG_ numbers held steady over time in the livers of liver tumor-bearing mice (**Fig. 1B**).

**Figure 1.**
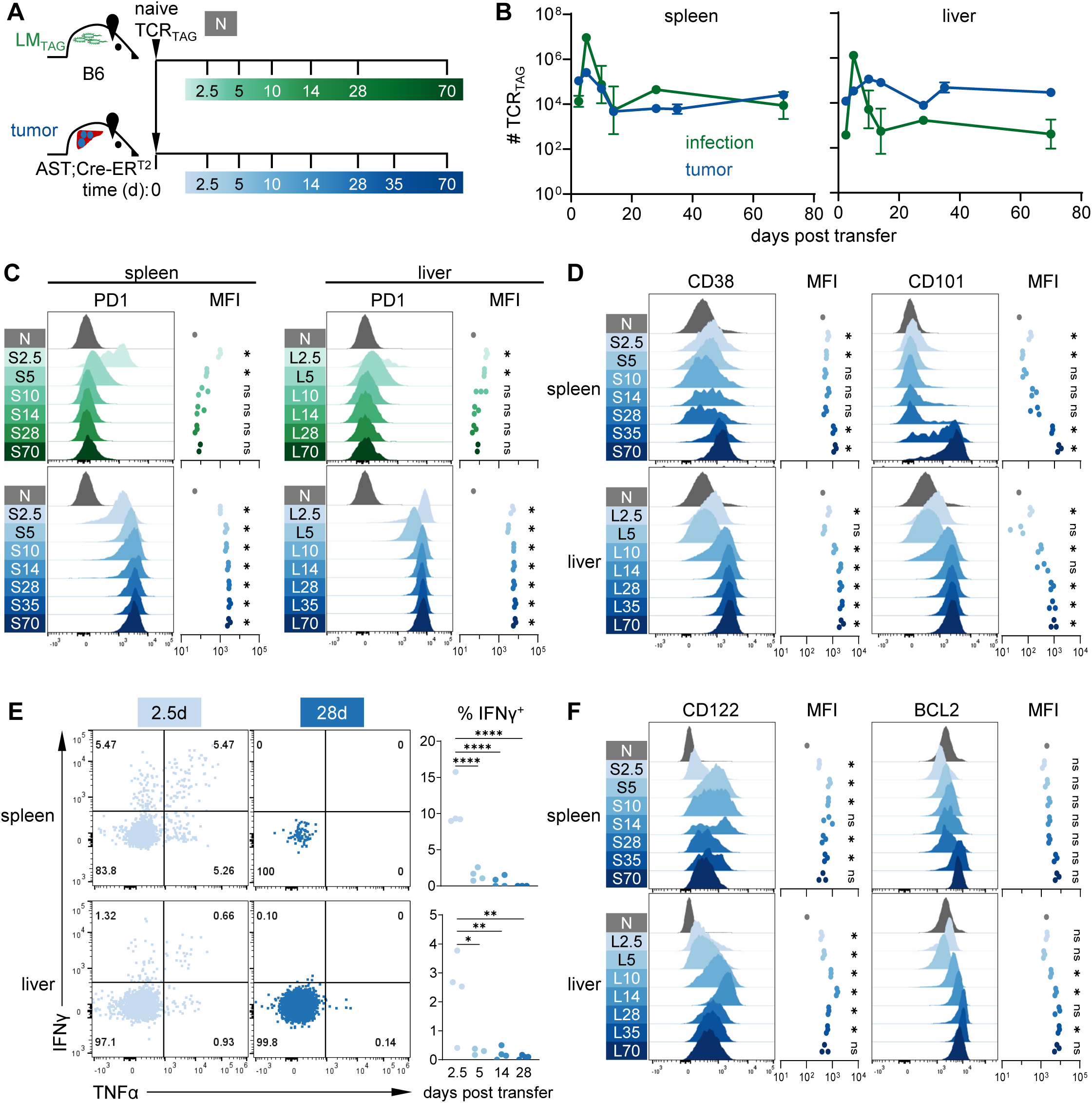
Tumor-specific CD8 T cell persist throughout tumorigenesis. **A.** Experimental scheme: naive TCR_TAG_ (Thy1.1) were adoptively transferred into LM_TAG_-infected C57BL/6 (B6; Thy1.2) or tamoxifen-treated AST;Cre-ER^T2^ (Thy1.2) mice. TCR_TAG_ were re-isolated from spleens and livers of infected mice (green) and tumor mice (blue) for flow cytometric analysis between 2.5-70+ days post-transfer. **B.** TCR_TAG_ cell numbers in the spleens (left) and livers (right) of LM_TAG_-infected mice and tumor-bearing mice. Dots represent mean and error bars represent SEM. For **B-D** and **F** n = 2-3 mice per group (infection); n = 3 mice per group (tumor). Data is representative of two independent experiments. **C.** Histograms (left) showing TCR_TAG_ PD1 expression from infection shown in comparison to naive (N; gray). For **C-D** and **F**, each symbol represents an individual mouse, and one-sample Student’s *t*-test with Bonferroni correction was used to determine significance: *, *P*<0.008 (infection); *, *P*<0.007 (tumor); ns, not statistically significant. **D.** Histograms and summary plots of CD38 expression and CD101 expression in spleens and livers from tumor mice shown in comparison to N. **E.** TCR_TAG_ IFNγ and TNFα production after 4-hour *ex vivo* TAG peptide stimulation, with inset numbers indicating the percentage of cells in each gate. Gates were set based on no peptide stimulation controls. Right, summary plots of % IFNγ^+^ TCR_TAG_ after peptide stimulation; each symbol represents an individual mouse; n = 3-4 mice per group. *, *P*<0.05; **, *P*<0.01; ****, *P*<0.0001 (one-way ANOVA with post hoc Tukey test). **F.** Histograms and summary plots of CD122 expression and BCL2 expression from tumor mice shown in comparison to N. All flow plots are gated on live CD8^+^ Thy1.1^+^ cells, and flow data for each timepoint is concatenated from all biological replicates. Summary plots (right) show MFI (geometric <u>m</u>ean fluorescence intensity).

TCR_TAG_ in infected mice showed transient upregulation of the early activation/inhibitory marker PD1, while TCR_TAG_ in AST;Cre-ER^T2^ mice showed persistent high-level PD1 expression in both the spleen and liver (**Fig. 1C**). We previously demonstrated that the expression of cell surface proteins such as CD38 and CD101 is associated with fixed-dysfunctional epigenetic states (10). Interestingly, while CD38 and CD101 were upregulated in liver-infiltrating TST 10-14 days after transfer, TST in the spleen did not upregulate these late-dysfunctional markers until day 28 and beyond (**Fig. 1D**), suggesting that dysfunction progresses more rapidly within tumors due to higher antigen exposure. At early time points (2.5-5 days), a subset of TST in AST;Cre-ER^T2^ spleens and a smaller subset in livers produced effector cytokines TNFα or IFNγ in response to TAG peptide stimulation; this subset rapidly diminished over time (**Fig. 1E**). In contrast, TCR_TAG_ in LM_TAG_-infected mice showed robust effector cytokine production capacity early after transfer and at memory time points, following the classical effector and memory differentiation trajectory in acute infection (**Supplementary Fig. 1B**).

We profiled the expression of activation/homing markers, inhibitory receptors, cytokine receptors, proliferation and survival markers, and transcription factors regulating T cell differentiation in infection and tumors (summary heatmaps shown in **Supplementary Fig. 2A**, histogram plots shown in **Supplementary Fig. 2B, C**). TST downregulated lymphoid homing receptor CD62L and progressively upregulated CD69, a marker of early activation but also a mediator of tissue residency through S1PR1 antagonism (24). The inflammatory cytokine IL2 (25)and homeostatic cytokines IL7 (26)and IL15 (27) play a crucial role in the proliferation and survival of CD8 T cells (reviewed in (28)). While early effector (day 2.5) TCR_TAG_ in LM_TAG_-infected hosts upregulated CD25/IL2Rα upon activation, TST failed to do so (**Supplementary Fig. 2A, C**). TST downregulated CD127/IL7Rα, while infection-activated TCR_TAG_ progressively upregulated CD127/IL7Rα (**Supplementary Fig. 2A, C**). However, both infection-activated T cells and TST upregulated and maintained CD122/IL2Rβ expression (**Fig. 1F**, **Supplementary Fig. 2A, C**), a subunit of the IL2 and IL15 receptor complex (29) consistent with previous work showing that IL15 signaling is critical for TST survival in tumors (30). The anti-apoptotic molecule BCL2 is required for cytokine-mediated survival (31) and expressed in long-lived memory T cells (32). Accordingly, TST upregulated BCL2 early and maintained its expression throughout tumorigenesis (**Fig. 1F, Supplementary Fig. 2A, C**). TST rapidly upregulated TOX, a transcription factor shown to be required for long-term survival in tumors (33) and during chronic viral infection (34–36), in both the spleen and liver (**Supplementary Fig. 2A, C**). Together, these data demonstrate that TST persist throughout tumorigenesis, expressing memory-associated pro-survival molecules.

### A subset of TCF1^lo^ TST actively cycle in the liver throughout tumorigenesis

To determine whether the TST population was maintained by continued proliferation or by long-lived post-mitotic cells, we performed cell cycle analysis on TST across time points from 2.5-70 days post-transfer (**Fig. 2A**). Following transfer and upon activation, most TST entered cell cycle (G_1_), as shown by KI67 upregulation in both the spleens and livers of tumor-bearing mice (**Fig. 2B**). We found that early after transfer, up to 30% of TST in the liver were actively cycling (S-G_2_/M) (**Fig. 2C**). Subsequently, TST began exiting cell cycle, as shown by KI67 downregulation, (**Fig. 2B**), with a progressive decrease in the number of TST actively cycling (**Fig. 2C**). Interestingly, from day 14 on, there was a small but stable subset of actively cycling TST in the liver (**Fig. 2C**), raising the question whether cycling TST within liver tumors were a TCF1^hi^ stem-like/progenitor population that could sustain the TST population long-term. Following transfer into tumor-bearing hosts, TST in the spleen initially expressed TCF1. However, from day 14 on, TST downregulated TCF1, and by day 35 most TST in the spleen were TCF1^lo^ (**Fig. 2D**). A similar transition occurred earlier in liver TST, which were largely TCF1^lo^ from day 10 on (**Fig. 2D**). Surprisingly, even as the number of TCF1^hi^ TST declined precipitously in the spleen (day 28 on) and liver (day 10 on), the total number of TST in the spleen and liver remained stable (**Fig. 2E**), and it was in fact TCF1^lo^ TST that comprised the majority of cycling TST in the liver from day 10 on (**Fig. 2F**).

**Figure 2.**
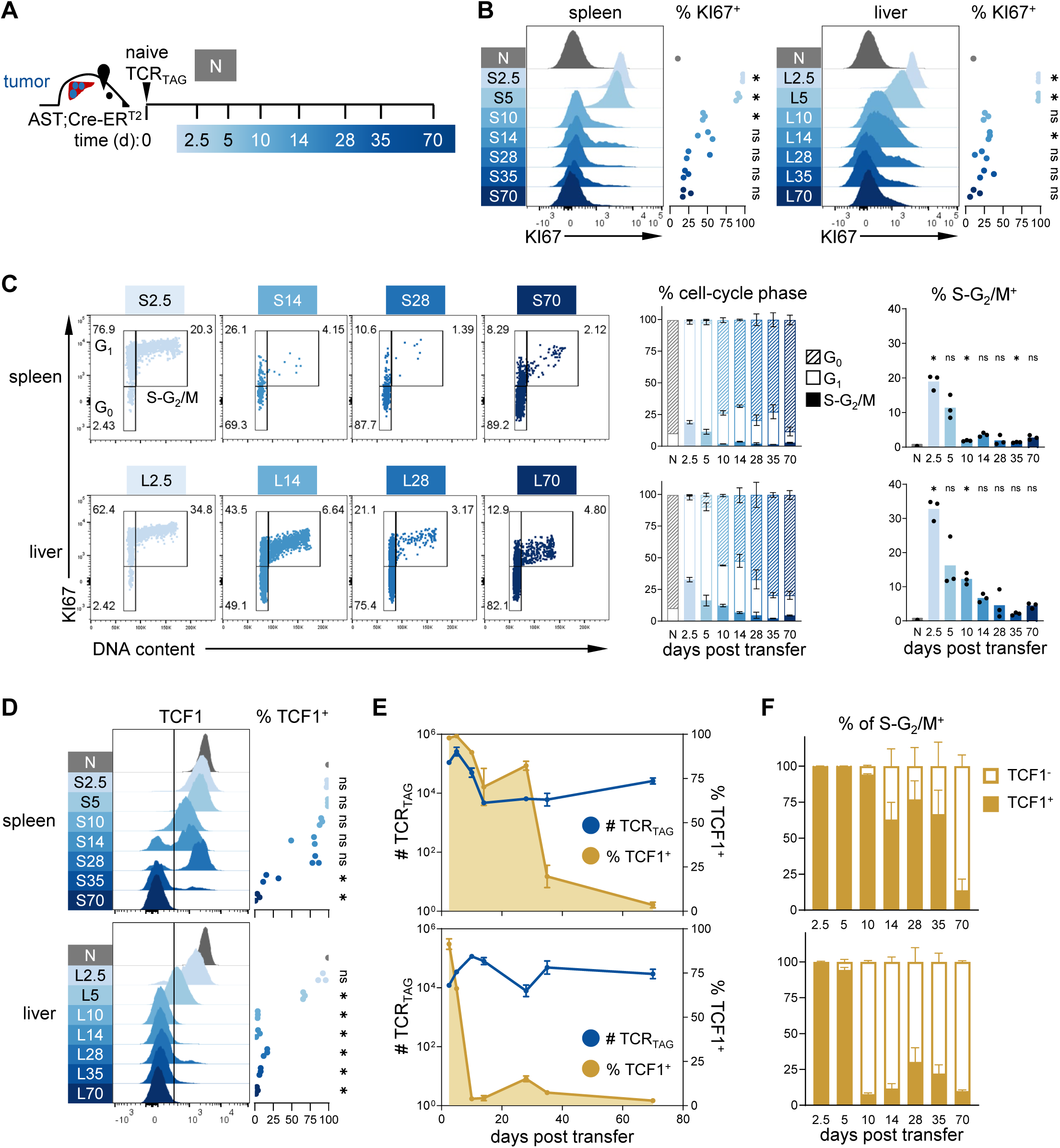
A subset of TCF1^lo^ TST actively cycle in the liver throughout tumorigenesis. **A.** Experimental scheme: naive TCR_TAG_ (Thy1.1) were adoptively transferred into tamoxifen-treated AST;Cre-ER^T2^ (Thy1.2) mice. TCR_TAG_ were re-isolated from spleens and livers for flow cytometric analysis at 2.5, 5, 10, 14, 28, 35, and 70+ days post-transfer. **B.** Histograms and summary plots of KI67 expression of TCR_TAG_ in spleens and livers from tumor-bearing mice shown in comparison to N, with % KI67^+^ gate set to exclude N. Flow plots for each time point were concatenated from all biological replicates; each symbol represents an individual mouse. *, *P*<0.007 determined by one-sample Student’s *t*-test with Bonferroni correction. **C.** Left, representative dot plots of KI67 expression and DNA content staining in TCR_TAG_ in spleens and livers of AST;Cre-ER^T2^ mice, with inset numbers indicating the percentage of cells in G_0_, G_1_, and S-G_2_M phases of cell cycle. Middle, summary plots of the percentage of TCR_TAG_ in G_0_, G_1_, and S-G_2_M, shown as mean ± SEM. Right, summary plots of the percentage of TCR_TAG_ in S-G_2_M. Each symbol represents an individual mouse. *, *P*<0.007 determined by one-sample Student’s *t*-test with Bonferroni correction. **D.** Histograms and summary plots of TCF1 expression in spleens and livers, with TCF1^+^ gate shown. Flow plots for each time point were concatenated from all biological replicates; each symbol represents an individual mouse. *, *P*<0.007 determined by one-sample Student’s *t*-test with Bonferroni correction. **E.** Longitudinal analysis of the absolute cell numbers of TCR_TAG_ cells (blue) in the spleens and livers of tumor-bearing mice and the % TCF1^+^ TCR_TAG_ (yellow), shown as mean ± SEM. **F.** Summary plots of the % TCF1^+^ (filled bars) and TCF1^−^ (open bars) TCR_TAG_ within the S-G_2_/M^+^ subset, shown as mean ± SEM. Data is representative of two independent experiments with n = 3 mice per group

### TST undergo stochastic proliferation and exit cell cycle

To determine whether a progenitor population maintains the TST population in tumors, we utilized dual-labeling with 5-ethynyl-2’-deoxyuridine (EdU) and 5-bromo-2’-deoxyuridine (BrdU), thymidine analogs incorporated into DNA during S phase. Naive TCR_TAG_ were adoptively transferred into tamoxifen-treated AST;Cre-ER^T2^ mice or LM_TAG_-infected B6 mice; an EdU pulse was administered 35 days post-transfer, followed by a BrdU pulse 7 days later (**Fig. 3A**). Progenitor cells that continuously self-renew would be dual-labeled with EdU and BrdU (**Fig. 3B left**), as observed in multipotent progenitors during hematopoiesis (37). In contrast, if TST enter and exit cell cycle in a stochastic manner, proliferated cells would be single-labeled with either EdU or BrdU, with few dual-labeled cells (**Fig. 3B right**). We first examined memory TCR_TAG_ generated after LM_TAG_ infection and found only single-labeled EdU^+^ or BrdU^+^ (**Fig. 3C**), consistent with previous work showing that memory T cells undergo stochastic homeostatic proliferation (38). Interestingly, we found few or no dual-labeled TST in liver tumors; rather, proliferated TST had incorporated either EdU or BrdU (**Fig. 3C**), suggesting that TST enter and exit cell cycle in a stochastic manner. Moreover, nucleoside-labeled TST were TCF1^lo^, in contrast to labeled memory TCR_TAG_ which were TCF1^hi^ (**Fig. 3D**), further evidence that TCF1 is not required for sustained TST proliferation in tumors.

**Figure 3.**
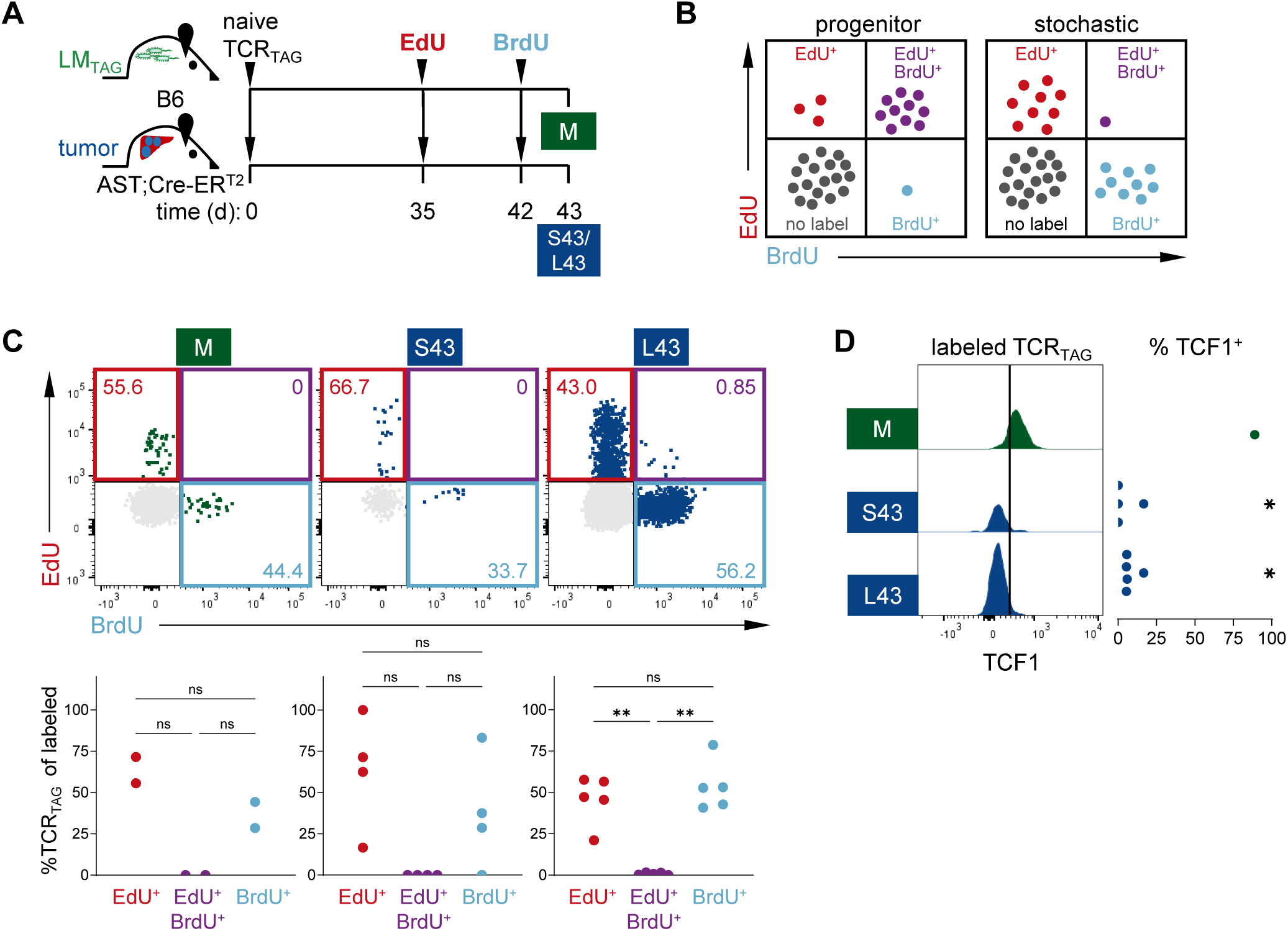
TST undergo stochastic proliferation and exit cell cycle. **A.** Experimental scheme: naive TCR_TAG_ (Thy1.1) were adoptively transferred into LM_TAG_-infected B6 (Thy1.2) or tamoxifen-treated AST;Cre-ER^T2^ (Thy1.2) mice and 5-ethynyl-2’-deoxyurdine (EdU) and 5-bromo-2’-deoxyuridine (BrdU) were administered on days 35 and 42, respectively. TCR_TAG_ were re-isolated on day 43 from spleens of infected mice (green) and spleens and livers of tumor-bearing mice (blue) for flow cytometric analysis. **B.** Schematic representation of the expected distributions of EdU and BrdU incorporation for T cells proliferating in a progenitor-progeny hierarchy (left) or stochastically (right). **C.** Top, dot plots of TCR_TAG_ EdU and BrdU incorporation in infected mice and tumor-bearing mice. Unlabeled cells shown in gray with inset numbers showing the percentage of nucleoside labeled cells in each gate. Bottom, summary plots of % EdU^+^ (red), EdU^+^,BrdU^+^ (purple), and BrdU^+^ (cyan) TCR_TAG_ within each nucleoside labeled subset. Each symbol represents an individual mouse. **, *P*<0.01 determined by repeated measures, one-way ANOVA with post hoc Tukey test. **D.** Concatenated histogram of TCF1 expression in nucleoside-labeled TCR_TAG_ and summary plot of the percentage of nucleoside-labeled TCR_TAG_ expressing TCF1. Each symbol represents an individual mouse. *, *P*<0.025 determined by one-sample Student’s *t*-test with Bonferroni correction with n=4-5 mice per group and representative of two independent experiments. Flow plots show data concatenated from all biological replicates.

### TCF1-knockout TST persist and proliferate in tumors

To directly test the requirement for TCF1 in TST stochastic proliferation/maintenance, we utilized CRISPR/Cas9-mediated *Tcf7* disruption. Naive TCR_TAG_;Cas9 CD8 T cells were transfected with a guide RNA (sgRNA) targeting *Tcf7* (TCF1KO) or a non-targeting control (NTC), leading to TCF1 protein downregulation (**Fig. 4A)**. TCF1KO or NTC TCR_TAG_ were adoptively transferred into tamoxifen-treated AST;Cre-ER^T2^ mice; 32 days later, mice were pulsed with EdU for 3 days prior to endpoint (**Fig. 4B**). Strikingly, TCF1KO TCR_TAG_ were undiminished in number in the spleens or livers of AST;Cre-ER^T2^ mice as compared to NTC TCR_TAG_ (**Fig. 4C**). At day 35, most TST proliferation (EdU^+^/KI67^+^) occurred in the liver (**Fig. 4D, E**), and TCF1KO showed robust EdU incorporation and at a similar frequency to NTC TCR_TAG_ (**Fig. 4D, E**). Thus, TCF1 does not appear to be necessary for TST stochastic proliferation and persistence in tumor-bearing hosts.

**Figure 4.**
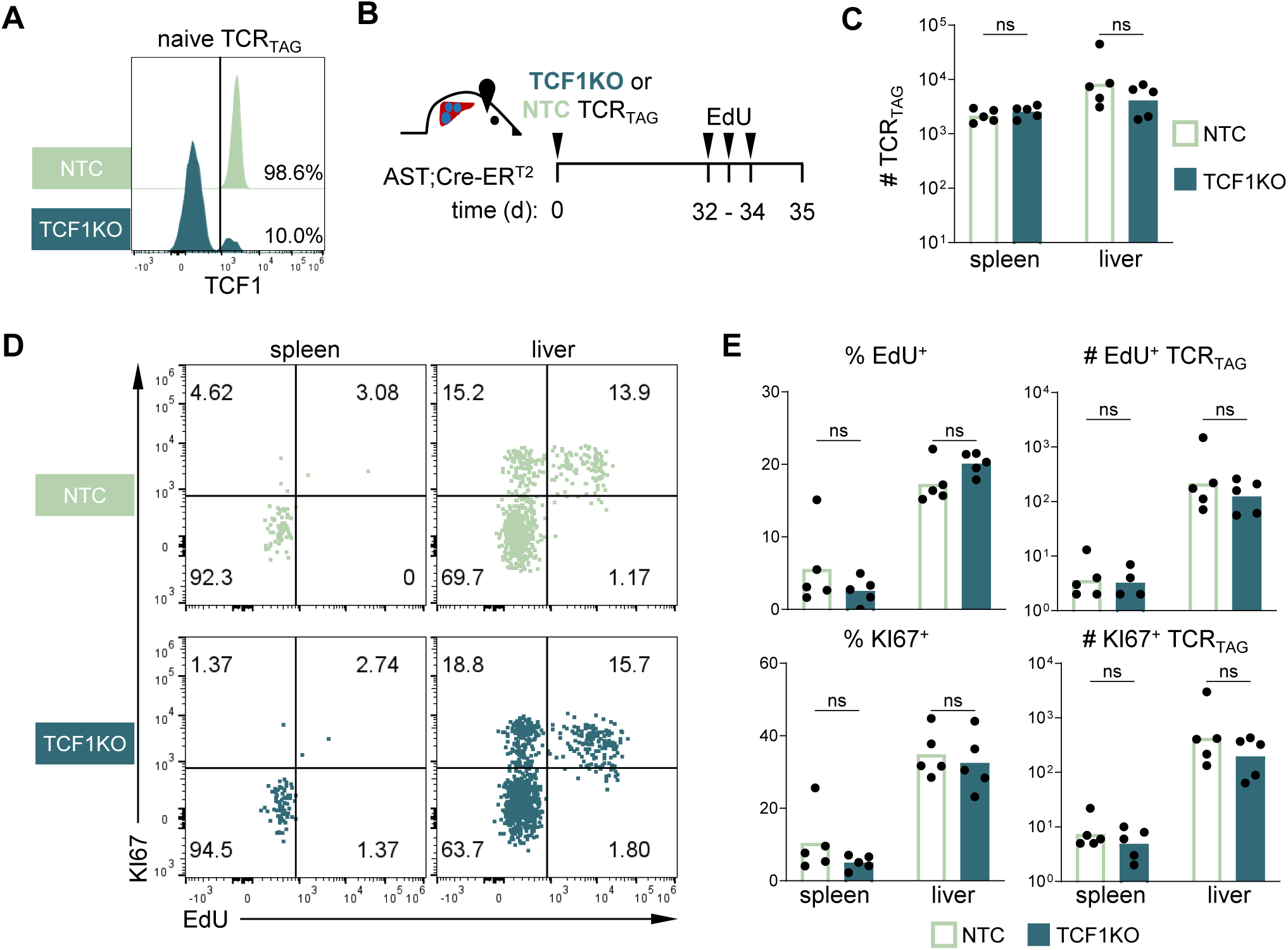
TCF1-knockout TST persist and proliferate in tumors. **A.** Histogram of TCF1 expression 48 hours post transfection of naive Cas9;TCR_TAG_ with sgRNA transfected with non-targeting control sgRNA (NTC; light green) or TCF7 sgRNA (TCF1KO; dark teal), data representative of two independent experiments. **B.** Experimental scheme: NTC or TCF1KO TCR_TAG_ (Thy1.1) were adoptively transferred into tamoxifen-treated AST;Cre-ER^T2^ (Thy1.2) mice. EdU was administered in the final 3 days. TCR_TAG_ were re-isolated on day 35 from the spleens and livers for flow cytometric analysis. **C.** NTC and TCF1KO TCR_TAG_ cell numbers recovered from tumor-bearing mice. **D.** Representative dot plots of KI67 expression and EdU incorporation of NTC and TCF1KO TCR_TAG_ cells, with gates set based on KI67^−^ /EdU^−^ cells. **E.** Top, summary plots of the percentage (left) and absolute number (right) of EdU^+^ NTC or TCF1KO TCR_TAG_. Bottom, summary plots of the percentage (left) and absolute number (right) of KI67^+^ or TCF1KO TCR_TAG_. For **C** and **E**, each symbol represents an individual mouse, with two-way ANOVA followed by post hoc Šídák’s multiple comparisons test with n=4-5 mice per group and representative of two independent experiments.

### TST persist in tumors without new influx from secondary lymphoid organs

We next asked whether the continued influx of TST from SLO is required to maintain the TST population in tumors, utilizing FTY720, a sphingosine-1-phosphate receptor agonist that inhibits lymphocyte egress from SLO. We adoptively transferred naive TCR_TAG_ into tamoxifen-treated AST;CreER^T2^ mice, with half of the mice treated with FTY720 or vehicle one day prior to T cell transfer and continuing for 5 days, and the other half treated from day 28 until endpoint at day 49 (**Fig. 5A**). All mice were pulsed with EdU for 3 days prior to endpoint. Within 24 hours of starting FTY720, few CD3ε-expressing T cells were found in peripheral blood as compared to vehicle (**Fig. 5B**), demonstrating effective inhibition of lymphoid egress from SLO. AST;Cre-ER^T2^ mice treated with FTY720 at the time of adoptive transfer had fewer TCR_TAG_ in the liver at day 5 (**Fig. 5C**), consistent with decreased early migration of TCR_TAG_ activated in SLO to the liver. Notably, FTY720 treatment initiated later (day 28) after the initial activation and contraction phase, did not impact the number of TCR_TAG_ in liver tumors (**Fig. 5C**). Moreover, TST EdU incorporation was similar in the livers of FTY720- and vehicle-treated mice (**Fig. 5D**), suggesting that liver-resident TST are the main proliferative population. EdU^+^ TST re-isolated from FTY720- and vehicle-treated AST;Cre-ER^T2^ mice after 5 days were TCF1^hi^, while EdU^+^ TST reisolated after 49 days were largely TCF1^lo^ (**Fig. 5E**). Most of the day 5 EdU^+^ TST expressed KI67 (**Fig. 5E**), consistent with an early activated and continuously cycling population(39). In contrast, day 49 EdU^+^ TST were largely negative for KI67 (**Fig. 5E**), consistent with cell cycle exit following EdU^+^. Thus, TCF1^lo^ TST in the liver enter and exit cell cycle, maintaining a stable population without new TST influx from SLO.

**Figure 5.**
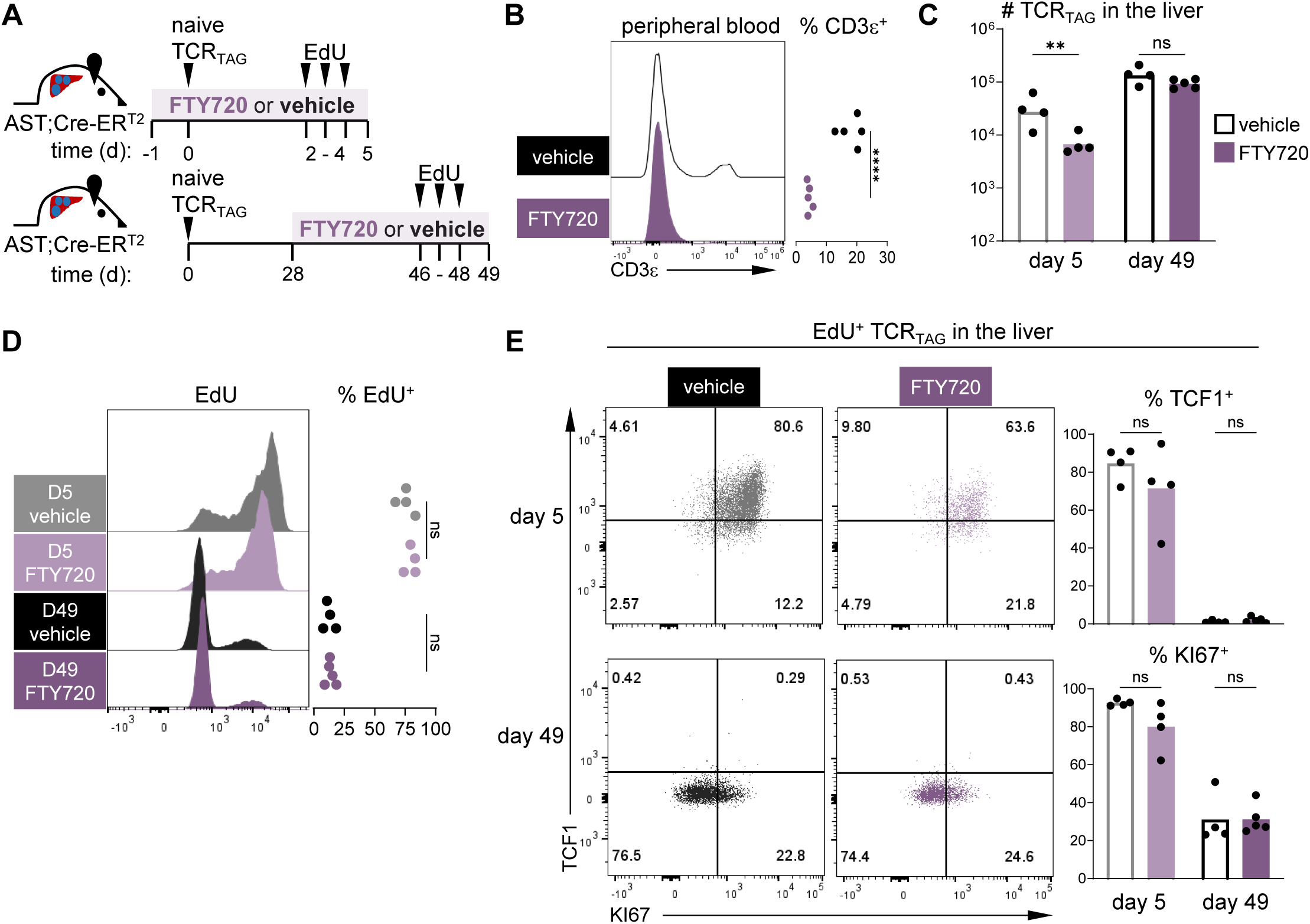
TST persist in tumors without new influx from secondary lymphoid organs. **A.** Experimental scheme: naive TCR_TAG_ (Thy1.1) were adoptively transferred into tamoxifen-treated AST;Cre-ER^T2^ (Thy1.2) mice. FTY720 treatment or vehicle was administered every other day beginning on day −1 for 6 days or day 28 for 3 weeks. EdU was administered for the final 3 days prior to harvest. TCR_TAG_ were re-isolated on day 5 and day 49 from the livers for flow cytometric analysis. **B.** Representative histogram and summary plot of % CD3ε^+^ cells in peripheral blood 24 hours after initiation of FTY720 (purple) or vehicle (black) treatment. ****, *P*<000.1 determined by unpaired, two-tailed Student’s *t*-test. **C.** TCR_TAG_ cell numbers in the livers of vehicle- (open black bars) and FTY720 (purple bars)-treated mice. Each symbol represents an individual mouse, with ****, *P*<0.0001 determined by two-way ANOVA followed by post hoc Šídák’s multiple comparisons test. **D.** Representative histogram of EdU+ incorporation in TCR_TAG_ T cells and summary plot of % EdU^+^ from the livers of vehicle- and FTY720-treated AST;Cre-ER^T2^ mice. Each symbol represents an individual mouse, two-way ANOVA followed by post hoc Šídák’s multiple comparisons test. **E.** Left, representative dot plots of TCF1 and KI67 expression in EdU^+^ TCR_TAG_ from the livers of AST;CreER^T2^ mice at day 5 (top) and day 49 (bottom) following vehicle and FTY720 treatment. Right, summary plots of % TCF1^hi^ (top) and KI67^+^ (bottom) of EdU^+^ TCR_TAG_. Each symbol represents an individual mouse, two-way ANOVA followed by post hoc Šídák’s multiple comparisons test. For **B-E**, n=4-5 mice per group and representative of two independent experiments.

To determine whether TST in other tumor histologies persist autonomously, we utilized a murine melanoma model. B16 murine melanoma cells expressing the CD8 T cell-recognized ovalbumin_257–264_ epitope (B16-OVA) were injected subcutaneously in B6 mice, treated with FTY720 starting on day 13 post-implantation, and pulsed with EdU during the final 3 days prior to endpoint (**Supplementary Fig. 3A**). FTY720 treatment led to loss of CD3ε^+^ T cells in the peripheral blood (**Supplementary Fig. 3B**). Endogenous, polyclonal OVA-specific CD8 T cells were identified using K^b^-OVA tetramer (OVA-TET). FTY720 treatment did not decrease the number of PD1^+^ OVA-TET^+^ CD8 T cells in tumors (**Supplementary Fig. 3C**). Moreover, EdU incorporation of OVA-specific CD8 T cells in the tumors was unaffected by FTY720 treatment; additionally, most tumor-resident T cells were TCF1^lo^ (**Supplementary Fig. 3D, E**). Thus, our studies demonstrate that following initial TST activation, expansion, and tumor infiltration, the TCF1^lo^ TST population is maintained autonomously through stochastic proliferation, without requiring additional TST influx from SLO.

## DISCUSSION

In this study, we leveraged our clinically relevant autochthonous liver cancer model to analyze TST dynamics throughout tumorigenesis and found that TCF1^lo^ TST were present in tumors and SLO throughout tumorigenesis and stably maintained for months. These studies are not feasible in patients with cancer, and transplantable murine cancer models do not permit such long-term assessment of TST. Thus, our study provides a unique and in-depth longitudinal analysis of TST during tumorigenesis and provides novel insights into how expression of activation/homing receptors, inhibitory receptors, cytokine receptors, functional markers, and transcription factors change in TST over time. While TST in tumor-bearing hosts were largely TCF1^hi^ at early time points following activation, within a few weeks, nearly all TST in the SLO and tumors were TCF1^lo^. Nevertheless, these TCF1^lo^ TST continued to proliferate, as shown by the ability to incorporate EdU/BrdU, even at late time points. Tumor-resident TST were able to proliferate and persist even after genetic deletion of TCF1 and when new influx of TST from SLO was blocked with FTY720.

While TCF1^hi^ T cells have been the focus and target for therapies as they are thought to mediate persistence and ICB responses, there is a great deal of heterogeneity in immune infiltration and T cell composition of tumors across tumor types and between patients. Moreover, the presence of TCF1^hi^ T cells in tumors does not always reliably predict better ICB responses (16), and we previously showed that TCF1^hi^ memory-like T cells reactive against antigens expressed in tumors and normal tissue (shared antigens) are profoundly dysfunctional and unable to respond to ICB (40). The finding that TCF1^lo^ TST persist and proliferate long-term raises the question of whether these T cells can contribute to anti-tumor immunity. Accordingly, a recent study demonstrated that TCF1 expression was more critical in poorly immunogenic tumors; however, in immunogenic tumors, TCF1-knockout TST responded to ICB with increased anti-tumor function (41).

Given that TCF1^lo^ TST proliferate and persist long-term and are found in robust numbers within tumors, the next step is to design novel strategies to target and reprogram these tumor-specific T cells within tumors. We showed that TCF1^lo^ TST express high levels of CD122 (IL2Rβ), a subunit of the IL2 and IL15 receptor complex (29), consistent with previous work showing that IL15 promotes the persistence of TCF1^lo^ tumor-infiltrating T cells (30). Numerous clinical trials are underway testing IL15-based therapeutics (reviewed in (42)), and it remains to be seen whether IL15 agonists are sufficient to reprogram TCF1^lo^ TST on their own or must be combined with other modalities. A recent study showed that a recombinant Fc-IL4 fusion protein promoted the survival of functional TCF1^lo^ TST and enhanced the efficacy of adoptive cell therapy and ICB (43). Proliferation/cell cycle is hypothesized to be the optimal window for cellular reprogramming (44,45). Thus, future studies to elucidate and target proliferation mechanisms in TCF1^lo^ TST could provide an opportunity for targeted TST epigenetic reprogramming during proliferation, restoring cytotoxicity and improving cancer immunotherapy.

## METHODS

### Mice

All mice were bred and maintained in a specific pathogen free barrier facility at Vanderbilt University Medical Center (VUMC, Nashville, TN). The mouse housing facility was on 12-hour light-dark cycles and maintained at 20–25 °C and 30–70% humidity. Mice were age- and sex-matched, between 6 and 12 weeks old when used for experiments and assigned randomly to experimental groups. Both female and male mice were used. TCR_TAG_ transgenic mice (B6.Cg-Tg(TcraY1,TcrbY1)416Tev/J, RRID:IMSR_JAX:005236) (46), Cre-ER^T2^ mice (B6.129-Gt(ROSA)26Sortm1(cre/ERT2)Tyj/J, RRID:IMSR_JAX:008463), Rosa26-Cas9 mice (Gt(ROSA)26Sortm1.1(CAG-cas9*,-EGFP)Fezh/J, RRID:IMSR_JAX:026179), B6 Thy1.1 mice (B6.PL-Thy1a/CyJ, RRID:IMSR_JAX:000406), and B6 mice (C57BL/6J, RRID:IMSR_JAX:000664) were purchased from the Jackson Laboratory. AST [AlbuminfloxStop-SV40 large T antigen (TAG)] mice, obtained from Natalio Garbi and Günter Hämmerling, German Cancer Research Center and previously described (47), were crossed to Cre-ER^T2^ mice to generate AST;Cre-ER^T2^. TCR_TAG_ mice were crossed to B6 Thy1.1 mice to generate TCR_TAG_;Thy1.1 mice. TCR_TAG_;Thy1.1 mice were crossed to Rosa26-Cas9 mice to generate Cas9; TCR_TAG_;Thy1.1/Thy1.2 mice. All animal experiments were performed in compliance with VUMC Institutional Animal Care and Use Committee (IACUC) regulations and in accordance with approved VUMC IACUC protocol M1700166-02.

### Listeria infection

The *Listeria monocytogenes* (*Lm*) *Δ actA Δ inlB* strain (48) expressing the TAG-I epitope (SAINNYAQKL, SV40 large T antigen_206-215_) (LM_TAG_) was generated by Aduro Biotech (22). Naive B6 mice were infected intravenously with 5 × 10^6^ colony forming units of LM_TAG_ 1 hour after T cell adoptive transfer

### Tamoxifen administration

Tamoxifen solution was prepared by warming tamoxifen (Sigma Aldrich, Cat# T5648, Cas-No: 10540-29-1) at 50°C for 1 hour in sterile corn oil (Sigma Aldrich, Cat# C8267, Cas-No: 8001-30-7) with 5% absolute ethanol (Sigma Aldrich, Cat# E7023, Cas-No: 67-15-5). A single dose of tamoxifen (1mg) was administered intraperitoneally in AST-CreER^T2^ mice 5 days prior to adoptive transfer.

### Adoptive transfer

To transfer naive TCR_TAG_ T cells into AST;CreER^T2^ or B6 mice, spleens from naive TCR_TAG_;Thy1.1 transgenic mice were mechanically disrupted with the back of 3 mL syringe and filtered through a 70 µm strainer into ammonium chloride potassium (ACK) buffer (150 mmol/L NH_4_Cl, 10 mmol/L KHCO_3_, 0.1 mmol/L Na_2_EDTA) to lyse erythrocytes. Cells were washed twice with cold serum-free RPMI 1640 (Corning, Cat# 10-040-CV) media and resuspended in serum-free RPMI for transfer. 2.5×10^6^ TCR_TAG_;Thy1.1 CD8^+^ T cells were adoptively transferred into induced AST;Cre-ER^T2^ mice. 5×10^5^ TCR_TAG_;Thy1.1 were adoptively transferred into B6 (Thy1.2) mice 1 hour prior to inoculation with LM_TAG_.

### Cell isolation for subsequent analyses

Spleens from experimental mice were mechanically disrupted with the back of 3 mL syringe and filtered through a 70 µm strainer into ACK buffer. Cells were washed once and resuspended in cold RPMI 1640 media supplemented with 10% fetal calf serum (FCS; Corning, Cat# 35-010-CV) (RPMI-10). Liver tissue was mechanically disrupted using a 150 µm metal mesh and glass pestle in cold phosphate buffered saline (PBS) containing 2% FBS (FBS/PBS) and passed through a 70 µm strainer. Liver homogenate was centrifuged at 400g for 5 min at 4°C and supernatant discarded. Liver pellet was resuspended in 15 mL of 2% FBS/PBS buffer containing 500 U heparin (NDC 63323-540-05), mixed with 10 mL of PBS-buffered Percoll (Cytiva, Cat# 17089101) by inversion, and centrifuged at 500g for 10 min at 4°C. Supernatant was discarded and pellet was RBC lysed in ACK buffer and resuspended in RPMI-10 for downstream applications.

### Antibodies and reagents

The following fluorochrome-conjugated antibodies were used in the analysis. The clone is denoted in parentheses.

Anti-BCL2-PE/Cyanine7 (BCL/10C4), BioLegend Cat# 633512, RRID:AB_2565247;

Anti-BIM-Alexa Fluor 488 (C34C5), Cell Signaling Technology Cat# 94805, RRID:AB_2800234;

Anti-BrdU-Alexa Fluor 647 (MoBu-1), Thermo Fisher Scientific Cat# B35133, RRID:AB_2536437;

Anti-CD3ε-PE (145-2C11), Tonbo Biosciences Cat# 50-0031, RRID:AB_2621730;

Anti-CD8a-PE/Cyanine7 (53-6.7), BioLegend Cat# 100722, RRID:AB_312761;

Anti-CD8a-Brilliant Violet 711 (53-6.7), BioLegend Cat# 100748, RRID:AB_2562100;

Anti-CD8a-APC/Cyanine7 (53-6.7), BioLegend Cat# 100714, RRID:AB_312753;

Anti-CD25-APC-eFluor780 (PC61.5), Thermo Fisher Scientific Cat# 47-0251-80, RRID:AB_1272215;

Anti-CD38-APC/Fire 750 (90), BioLegend Cat# 102738, RRID:AB_2876402;

Anti-CD39-PerCP-eFluor 710 (24DMS1) Thermo Fisher Scientific Cat# 46-0391-80, RRID:AB_10717513;

Anti-CD44-FITC (IM7), Tonbo Biosciences Cat# 35-0441, RRID:AB_2621688;

Anti-CD62L-Brilliant Violet 785 (MEL-14), BioLegend Cat# 104440, RRID:AB_2629685;

Anti-CD69-PE/Cyanine7 (H1.2F3), Thermo Fisher Scientific Cat# 25-0691-82, RRID:AB_469637;

Anti-CD90.1-Brilliant Violet 510 (OX-7), BioLegend Cat# 202535, RRID:AB_2562643;

Anti-CD90.1-PerCP/Cyanine5.5 (OX-7), BioLegend Cat# 202516, RRID:AB_961437;

Anti-CD90.1-APC (OX-7), BioLegend Cat# 202526, RRID:AB_1595470;

Anti-CD101-PE/Cyanine7 (Moushi101), Thermo Fisher Scientific Cat# 25-1011-80, RRID:AB_2573377;

Anti-CD122-PE (5H4), BioLegend Cat# 105905, RRID:AB_2125737;

Anti-CD127-PE/Cyanine7 (A7R34), BioLegend Cat# 135014, RRID:AB_1937265;

Anti-CD132-APC (TUGm2), BioLegend Cat# 132307, RRID:AB_10643575;

Anti-IFNγ-APC (XMG1.2), BioLegend Cat# 505810, RRID:AB_315404;

Anti-KI67-PerCP-Cyanine5.5 (16A8), BioLegend Cat# 350520, RRID:AB_2562295;

Anti-KI67-PE (16A8), BioLegend Cat# 652403, RRID:AB_2561524;

Anti-KI67-APC-eFluor 780 (SolA15), Thermo Fisher Scientific Cat# 47-5698-80, RRID:AB_2688064;

Anti-LEF1-PE (C12A5), Cell Signaling Technology Cat# 14440, RRID:AB_2798481;

Anti-Ly108-Pacific Blue (330-AJ), BioLegend Cat# 134608, RRID:AB_2188093;

Anti-PD1-PE/Cyanine (RMP1-30), BioLegend Cat# 109109, RRID:AB_572016;

Anti-PD1-APC (RMP1-30), BioLegend Cat# 109112, RRID:AB_10612938;

Anti-TCF1-PE (C63D9), Cell Signaling Technology Cat# 14456, RRID:AB_2798483;

Anti-TCF1-Alexa Fluor 647 (C63D9), Cell Signaling Technology Cat# 6709, RRID:AB_2797631;

Anti-Tim3-PE (RMT3-23), BioLegend Cat# 119703, RRID:AB_345377;

Anti-TNFα-PE (MP6-XT22), Thermo Fisher Scientific Cat# 12-7321-82, RRID:AB_466199;

Anti-TOX-APC (REA473), Miltenyi Biotec Cat# 130-118-474, RRID:AB_2751522.

The following cell dyes were used in the analysis.

4′,6-diamidino-2-phenylindole (DAPI), BioLegend, Cat# 422801, Cas-No: 47165-04-8;

Ghost Dye Violet 450 Viability Dye, Tonbo/Cytek, Cat# 13-0863;

Ghost Dye Red 710 Viability Dye, Tonbo/Cytek, Cat# 13-0871;

Ghost Dye Red 780 Viability Dye,Tonbo/Cytek, Cat# 13-0865.

APC-conjugated MHC class I (H2-K^b^) tetramer complexed with OVA_257-264_ peptide was prepared at the National Institutes of Health Tetramer Facility (Atlanta, GA).

### Cell surface and intracellular staining

Splenocytes or liver/tumor-infiltrating lymphocytes from naive TCR_TAG_ mice, *Listeria*-inoculated B6 mice, induced AST;Cre-ER^T2^ or B16-OVA inoculated B6 mice were stained with viability dye and antibodies against surface molecules in 2% FCS/PBS. The samples were then analyzed by flow cytometry (see Flow cytometric analysis). For intracellular staining, splenocytes or liver-infiltrating lymphocytes were surface stained as above and then fixed and permeabilized using the FoxP3 Transcription Factor Fix/Perm (Tonbo/Cytek, Cat# TNB-0607) per the manufacturer’s instructions before staining for intracellular molecules. The samples were then analyzed by flow cytometry. For analysis of effector cytokine production (IFNγ, TNFα), an ex vivo peptide stimulation was performed prior to staining. Splenocytes from naive, *Listeria*-inoculated, or induced AST;Cre-ER^T2^ mice were mixed with 2 × 10^6^ naive B6 splenocytes and incubated in RPMI-10 for 4 hours at 37°C in the presence of brefeldin A (Biolegend, Cat# 420601) and 0.5mmol/L of TAG epitope I peptide (Genscript, custom-synthesized). The cells were then surface-stained and fixed and permeabilized as described above and analyzed by flow cytometry.

### Flow cytometry

All flow analysis was performed on the Attune NXT Acoustic Focusing Cytometer (ThermoFisher Scientific). Data was analyzed using FlowJo v.10.10.0 (Tree Star Inc., RRID:SCR_008520).

### Cell cycle analysis

T cells were stained for surface and intracellular markers as described above (see Cell-surface and intracellular staining). Samples were suspended in FoxP3 Fix/Perm buffer (Tonbo/Cytek), and DAPI was added to samples immediately prior to flow cytometric analysis. Samples were gated to exclude “shadow doublets” (KI67^lo^ cells with > 2n DNA content) as previously described (49).

### EdU and BrdU labeling

5-ethynyl-2’-deoxyurdine (EdU) 3-day pulse labelling was performed by injecting 200µg/day of EdU (Sigma Aldrich, Cat# 900584, CAS-No: 61135-33-9) intraperitoneally. EdU incorporation was analyzed using the Click-iT Plus EdU Alexa Fluor 488 Flow Cytometry Assay Kit (ThermoFisher Scientific, Cat# C10633), according to manufacturer’s instructions, except FoxP3 Transcription Factor Fix/Perm (Tonbo/Cytek, Cat# TNB-0607) was used in place of provided fix/perm to allow for transcription factor staining. For dual labeling of EdU and 5-bromo-2’-deoxyuridine (BrdU; Sigma Aldrich, Cat# B50002, CAS-No: 59-14-3), 2mg of nucleoside analog was injected intraperitoneally. EdU Click-iT reaction was performed as described above prior to BrdU staining. Following EdU Click-iT reaction, cells were treated with DNase I (New England BioLabs Inc., Cat# M0303) for 10 minutes at 37°C followed by intracellular/transcription factor staining as described above, including the non-EdU cross-reactive anti-BrdU clone MoBU1 (50).

### FTY720 treatment

Fingolimod (FTY720) Hydrochloride (Selleckchem, Cat# S50002, CAS-No:162359-56-0) stock solution was prepared by dissolving FTY720 in dimethyl sulfoxide (DMSO; Sigma Aldrich, Cat# D8418, CAS-No: 67-68-5) to a concentration of 10mg/mL and stored at −80°C. FTY720 stock solution was diluted to 1 mg/mL in sterile 0.9% saline (NDC 0409-4888-02) immediately prior to administration. Sterile 0.9% saline containing DMSO (equivalent volume to FTY720 stock solution) was prepared as vehicle control. For AST;CreER^T2^ tumor studies, TCR_TAG_;Thy1.1 CD8^+^ T cells were transferred into mice as described above. One day prior to adoptive transfer, for day 5 time points, or 28 days after transfer, for day 49 timepoints, mice were treated by oral gavage with FTY720 (10 mg/kg) or vehicle control every other day until endpoint. For B16-OVA tumor studies, tumors were injected as described below. 13 days after tumor inoculation, mice were treated with FTY720 (10 mg/kg) or vehicle control intraperitoneally every other day for 8 days.

### B16-OVA tumor model

The B16 OVA_257-264_-Cerulean (B16-OVA) mouse melanoma cell line, expressing triple SIINFEKL-AAY repeats fused to Cerulean, was kindly provided by A. Schietinger (MSKCC) and previously described (51). The cell line tested negative for over 60 species of mycoplasma using the Universal Mycoplasma Detection Kit (ATCC, Cat# 30-1012K). B16-OVA cells were cultured in DMEM (Corning, Cat# 10-013-CV) supplemented with 10% FBS and penicillin-streptomycin-glutamine solution (cDMEM; Gibco, Cat# 10378-016) at 37°C in a 5% CO_2_ humidified incubator. For inoculation of subcutaneous tumors, B16-OVA was harvested at 60–80% confluency. Medium was replaced with fresh cDMEM the day before collection. Cells were washed twice with ice-cold serum-free RPMI and 1×10^6^ cells were injected subcutaneously into B6 mice. The maximal tumor size permitted for transplantable tumors as per our VUMC IACUC-approved protocol is 3 cm in maximum diameter. No tumors in this study exceeded the maximal allowed tumor size.

### T Cell isolation and electroporation/transfection

Spleens from naive Cas9;TCR_TAG_;Thy1.12 transgenic mice were mechanically disrupted with the back of 3 mL syringe and filtered through a 70 µm strainer into 2% FBS/PBS buffer. CD8+ T cells were magnetically isolated using EasySep^TM^ Mouse CD8+ T Cell Isolation Kit (STEMCELL Technologies, Cat# 19853A). T cells were then counted, pelleted, and resuspended in Opti-Mem (Gibco, Cat# 21985-023). A 100 μL suspension of T cells (7.5×10^6^ cells) and either sgRNA targeting *Tcf7* (GAAAGCUGGGGGACGCCAUG) (0.4nmol) (Genscript; custom-synthesized) or a non-targeting control sgRNA (GACAUUUCUUUCCCCACUGG) (0.4 nmol) (Genscript; custom-synthesized) were added to a 2mm sterile cuvette (Bulldog Bio, Cat# 12358-346). Cells were electroporated using NEPA21 Electro-Kinetic Transfection System (Bulldog Bio) with the following parameters: Poring Pulse: 250 V, 2 ms length, 50 ms interval, 2 pulses, 10% Decay Rate, + polarity; Transfer Pulse: 20 V, 50 ms length, 50 ms interval, 5 pulses, 40% Decay Rate, +/-Polarity. Immediately after electroporation, pre-warmed RPMI 1640 containing 10% FCS, penicillin-streptomycin-glutamine solution, 2-mercaptoethanol (Gibco, Cat# 21985-023), and 100ng/ml of recombinant human IL-7 (cRPMI; BRB Preclinical Biologics Repository) was added to the cuvette, and the cuvette was incubated at 37°C for 15 minutes. Cells were then gently mixed by pipetting and added to a flat-bottom 24-well plate. Additional warmed cRPMI was used to wash cuvette and added to wells. After 1-2 hours resting, cells were used for *in vivo* experiments.

### Statistical analyses and reproducibility

No statistical method was used to predetermine sample size, but sample sizes are similar to those reported in previous publications (17,40). For FTY720 treatment studies and TCF1KO studies, mice in AST;Cre-ERT2 tumor cohorts and B16-OVA tumor cohorts were randomly assigned to treatment groups. The investigators were not blinded to allocation during experiments and outcome assessment. All statistical analysis, including one-sample Student’s *t*-test with Bonferroni correction, one-way ANOVA with post hoc Tukey test, repeated measures, one-way ANOVA with post hoc Tukey test, two-way ANOVA with post hoc Šídák’s test, and unpaired, two-tailed Student’s *t*-test, were performed where indicated using GraphPad Prism v10 (RRID:SCR_002798). Data distribution was assumed to be normal, but this was not formally tested. Composite figures and schematics were generated using Microsoft PowerPoint 365 (RRID:SCR_023631).

### Data availability

The data generated in this study are available within the article and its Supplementary Data Files or are available from the corresponding author upon request.

## ACKNOWLEGEMENTS

We thank A. Schietinger, M. Boothby, D. McNitt, and members of the Philip laboratory for helpful discussions. We thank W. Bindeman for helpful discussions and technical assistance. We thank the Vanderbilt Division of Animal Care. We thank P. Lauer and Aduro Biotech for providing attenuated Listeria strains. We thank A. Schietinger and K.M. Hawley for technical assistance with the RNA electroporation protocol. This work was supported by the following funding sources: NIH T32AR059039 (M.M.E.), NIH T32GM008554 (N.R.F.), ASH Medical Student Physician-Scientist Award (L.A.B.), Medical Scientist Training Program (MSTP) NIH T32GM007347 to the Vanderbilt MSTP Program (M.W.R.), V Foundation Scholar Award (M.P.), NIH R37CA263614 (M.P.), Vanderbilt-Ingram Cancer Center (VICC) SPORE Career Enhancement Program (M.P.) NIH P50CA098131, Vanderbilt Digestive Disease Research Center (VDDRC) Young Investigator and Pilot Award (M.P.) NIH P30DK058404, and NIH T32CA009592 (C.R.D.R). The Vanderbilt Flow Cytometry Shared Resource (FCSR) maintained a group license for the FlowJo Software. The FCSR is supported by the Vanderbilt Ingram Cancer Center (P30 CA068485) and the Vanderbilt Digestive Disease Research Center (DK058404).

---

**Supplementary Figure 1.**
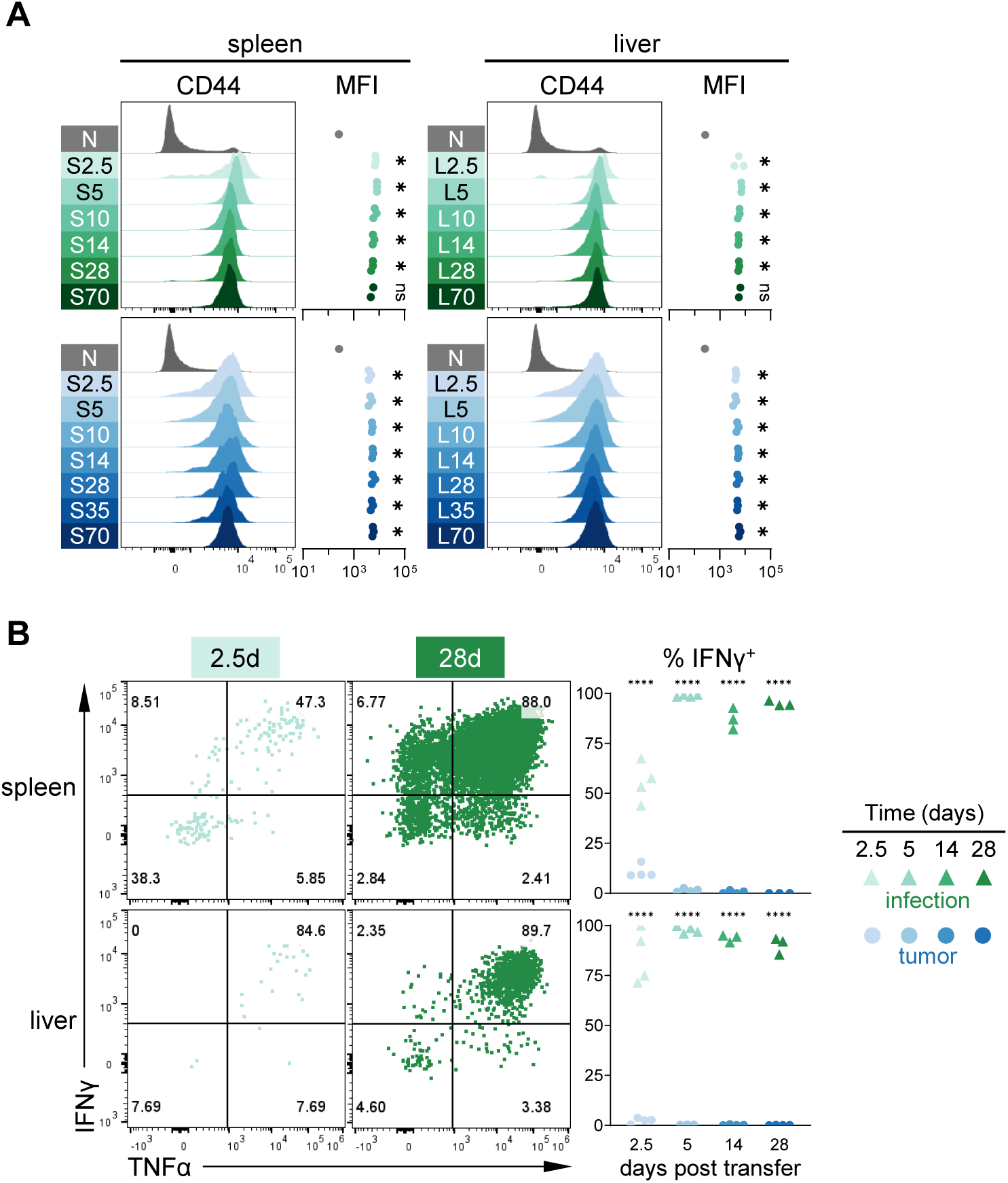
Antigen-specific CD8 T cell activation and cytokine production during acute infection and tumorigenesis. **A.** Experimental setup as shown in Fig. 1A. Histograms and summary plots of CD44 expression from infection (green) and tumor (blue) in spleen and liver shown in comparison to naive (N; gray). Each symbol represents an individual mouse, and one-sample Student’s *t*-test with Bonferroni correction was used to determine significance: *, *P*<0.008 (infection); *, *P*<0.007 (tumor); ns, not statistically significant. n = 2-3 mice per group (infection); n = 3 mice per group (tumor). Data is representative of two independent experiments. **B.** TCR_TAG_ IFNγ and TNFα production after 4-hour *ex vivo* TAG peptide stimulation, with inset numbers indicating the percentage of cells in each gate. Gates were set based on no peptide stimulation controls. Summary plots show % IFNγ^+^ TCR_TAG_ after peptide stimulation; each symbol represents an individual mouse; n = 3-4 mice per group. ****, *P*<0.0001 (two-way ANOVA followed by post hoc Šídák’s multiple comparisons test). All flow plots are gated on live CD8^+^ Thy1.1^+^ cells and flow data for each timepoint is concatenated from all biological replicates. Data is representative of two independent experiments.

**Supplementary Figure 2.**
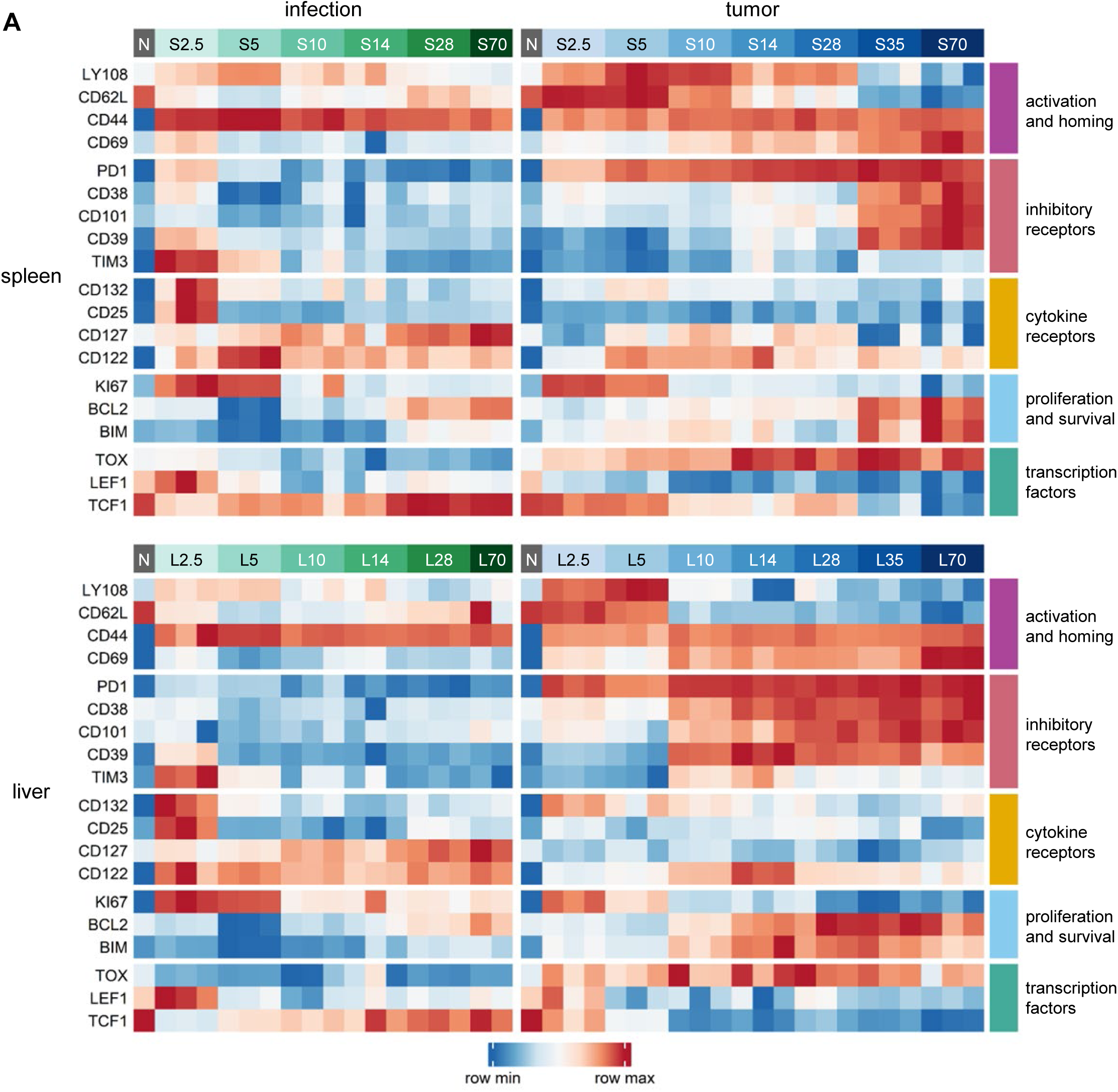
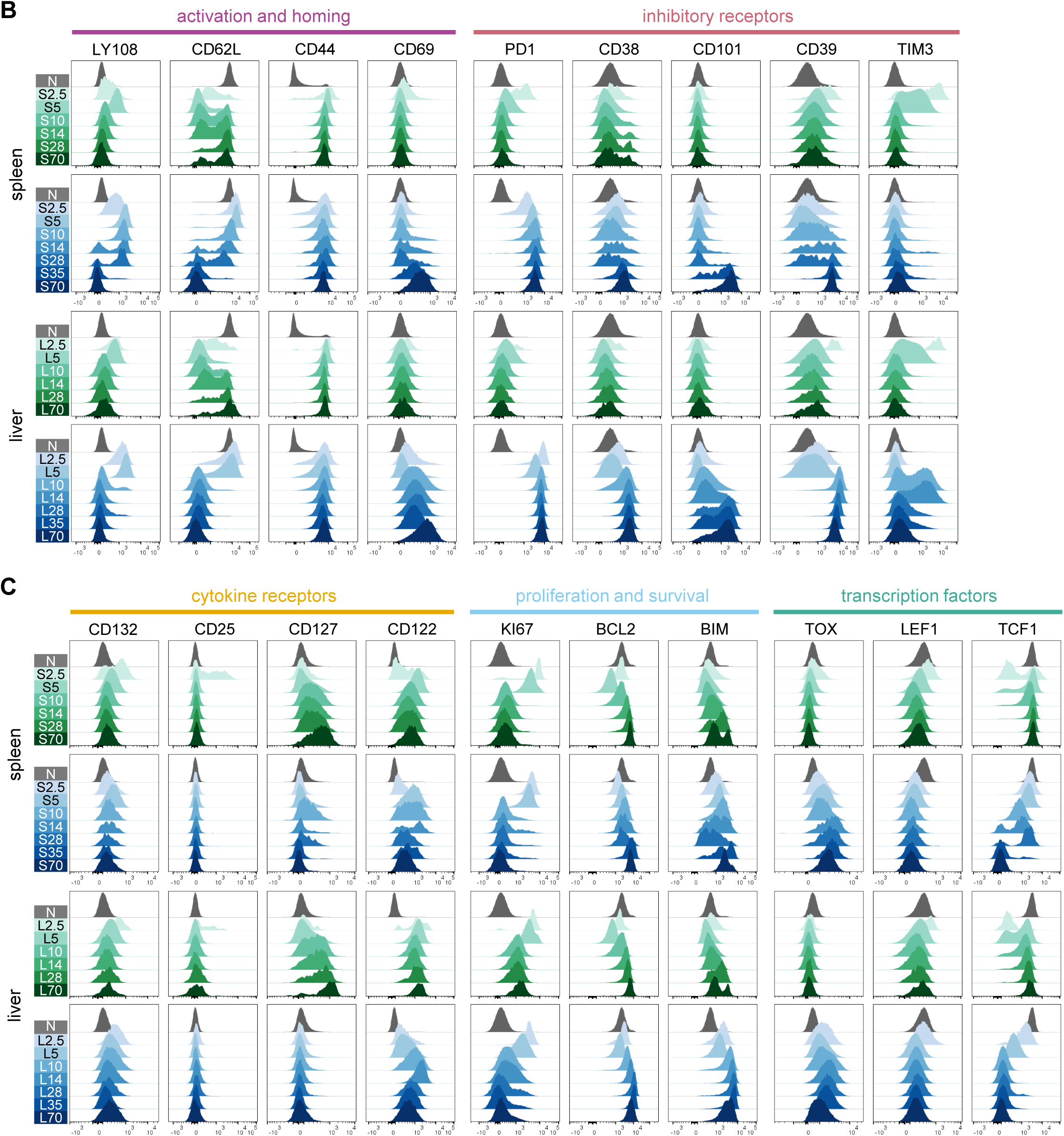
Antigen-specific CD8 T cell differentiation during acute infection and during tumorigenesis. **A.** Experimental setup as in Fig. 1A. Heatmap summarizing data for flow cytometric assessment of activation and homing proteins, inhibitory receptors, cytokine receptors, proliferation and survival markers, and transcription factors. Blue-red color scale shows row normalized log_2_ fold change (log2FC) of MFI in comparison to N. Each column represents individual biologic replicates for each condition and time point (the same single naive replicate is shown next to both infection and tumor groups). Data is representative of two independent experiments**. B.** Corresponding histograms showing expression of activation and homing markers (LY108, CD62L, CD44, CD69) andinhibitory receptors (PD1, CD38, CD101, CD101, and TIM3). **C.** Histograms showing expression of cytokine receptors (CD132, CD25, CD127, CD122), proliferation and survival markers (KI67, BCL2, BIM), and transcription factors (TOX, LEF1, and TCF1). All flow plots are gated on live CD8^+^ Thy1.1^+^ cells and flow data for each timepoint is concatenated from all biological replicates. Data is representative of two independent experiments.

**Supplementary Figure 3.**
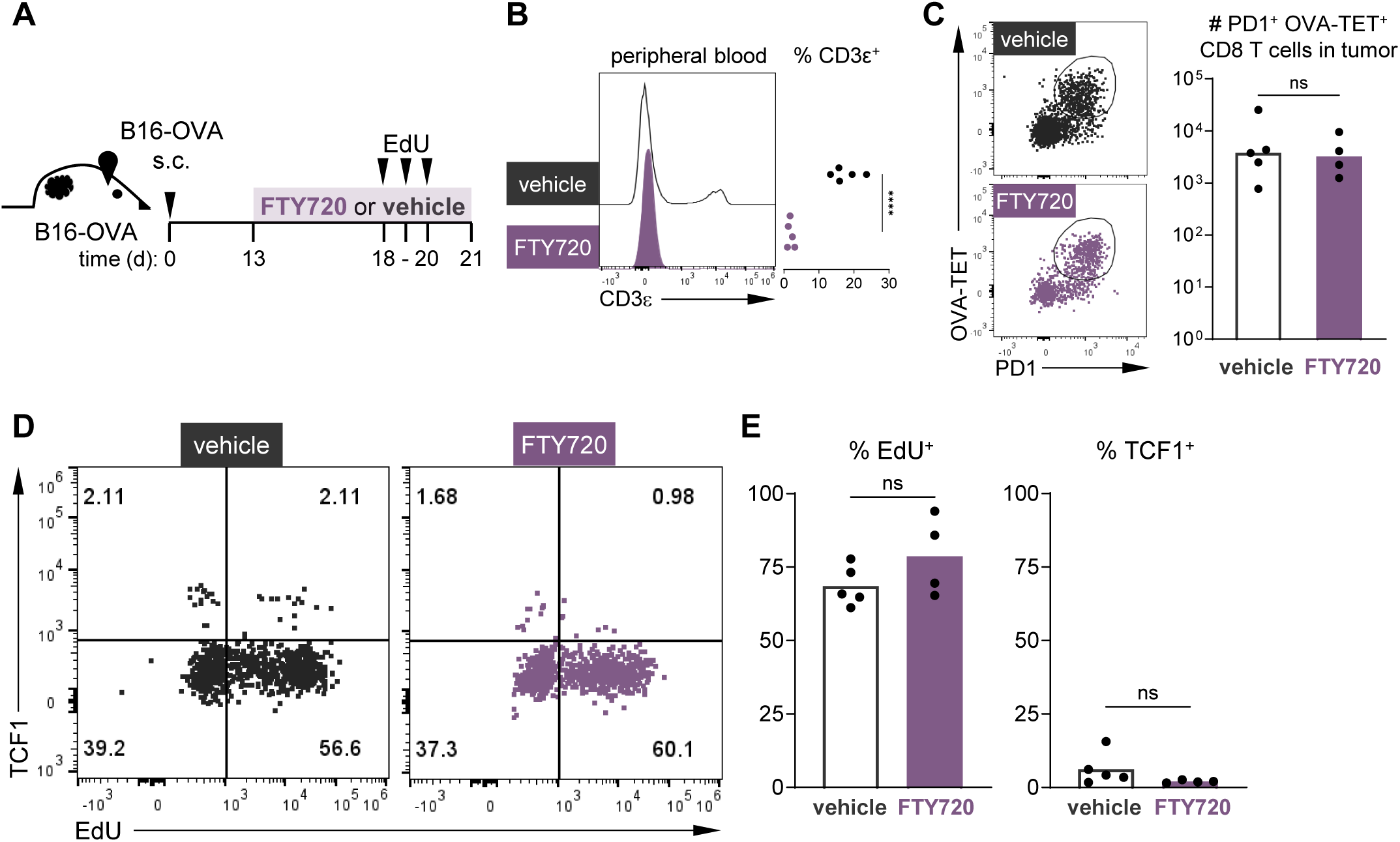
Endogenous TST in B16-OVA persist without new influx from secondary lymphoid organs. **A.** Experimental scheme: B16-OVA were injected subcutaneously into B6 mice. FTY720 treatment was administered every other day beginning on day 13 for 8 days. EdU was administered for the final 3 days. Lymphocytes were harvested from the tumors of mice on day 21 for flow cytometric analysis. **B.** Representative histogram and summary plot of CD3ε^+^ cells in peripheral blood 24 hours after initiation of FTY720 (purple) or vehicle (black) treatment. ****, *P*<000.1 determined by unpaired, two-tailed Student’s *t*-test. **C.** Representative dot plots showing gating of OVA tetramer^+^ (OVA-TET^+^)/PD1^+^ endogenous CD8 T cells and summary plot of OVA-TET^+^ PD1^+^ CD8^+^ T cell numbers recovered from vehicle- and FTY720-treated B16-OVA tumor-bearing mice. Each symbol represents an individual mouse, unpaired, two-tailed Student’s *t*-test. **D.** Representative dot plots of TCF1 expression and EdU incorporation in OVA-TET^+^ PD1^+^ CD8^+^ T cells from the tumors of vehicle- and FTY720-B16-OVA tumor-bearing mice. **E.** Summary plots of the percentage of EdU^+^ (left) and TCF1^hi^ (right) of OVA-TET^+^PD1^+^CD8^+^ T cells. Each symbol represents an individual mouse, unpaired, two-tailed Student’s *t*-test. For **B-E**, n=5 mice per group and representative of two independent experiments.

## REFERENCES

1. Haslam A, Olivier T, Prasad V. How many people in the US are eligible for and respond to checkpoint inhibitors: An empirical analysis. Int J Cancer 2025;156(12):2352–9 doi 10.1002/ijc.35347.

2. Mueller KP, Grenier JM, Weber EW. CAR T cell persistence in cancer. Trends Cancer 2025;11(10):1005–18 doi 10.1016/j.trecan.2025.08.014.

3. Wittling MC, Cole AC, Brammer B, Diatikar KG, Schmitt NC, Paulos CM. Strategies for Improving CAR T Cell Persistence in Solid Tumors. Cancers (Basel) 2024;16(16) doi 10.3390/cancers16162858.

4. Paley MA, Kroy DC, Odorizzi PM, Johnnidis JB, Dolfi DV, Barnett BE, et al. Progenitor and terminal subsets of CD8+ T cells cooperate to contain chronic viral infection. Science 2012;338(6111):1220–5 doi 10.1126/science.1229620.

5. Escobar G, Mangani D, Anderson AC. T cell factor 1: A master regulator of the T cell response in disease. Sci Immunol 2020;5(53) doi 10.1126/sciimmunol.abb9726.

6. He R, Hou S, Liu C, Zhang A, Bai Q, Han M, et al. Follicular CXCR5-expressing CD8(+) T cells curtail chronic viral infection. Nature 2016;537(7620):412–28 doi 10.1038/nature19317.

7. Im SJ, Hashimoto M, Gerner MY, Lee J, Kissick HT, Burger MC, et al. Defining CD8+ T cells that provide the proliferative burst after PD-1 therapy. Nature 2016;537(7620):417–21 doi 10.1038/nature19330.

8. Utzschneider DT, Charmoy M, Chennupati V, Pousse L, Ferreira DP, Calderon-Copete S, et al. T Cell Factor 1-Expressing Memory-like CD8(+) T Cells Sustain the Immune Response to Chronic Viral Infections. Immunity 2016;45(2):415–27 doi 10.1016/j.immuni.2016.07.021.

9. Philip M, Schietinger A. Heterogeneity and fate choice: T cell exhaustion in cancer and chronic infections. Curr Opin Immunol 2019;58:98–103 doi 10.1016/j.coi.2019.04.014.

10. Philip M, Fairchild L, Sun L, Horste EL, Camara S, Shakiba M, et al. Chromatin states define tumour-specific T cell dysfunction and reprogramming. Nature 2017;545(7655):452–6 doi 10.1038/nature22367.

11. Wu T, Ji Y, Moseman EA, Xu HC, Manglani M, Kirby M, et al. The TCF1-Bcl6 axis counteracts type I interferon to repress exhaustion and maintain T cell stemness. Sci Immunol 2016;1(6) doi 10.1126/sciimmunol.aai8593.

12. Gearty SV, Dundar F, Zumbo P, Espinosa-Carrasco G, Shakiba M, Sanchez-Rivera FJ, et al. An autoimmune stem-like CD8 T cell population drives type 1 diabetes. Nature 2022;602(7895):156–61 doi 10.1038/s41586-021-04248-x.

13. Sade-Feldman M, Yizhak K, Bjorgaard SL, Ray JP, de Boer CG, Jenkins RW, et al. Defining T Cell States Associated with Response to Checkpoint Immunotherapy in Melanoma. Cell 2018;175(4):998–1013 e20 doi 10.1016/j.cell.2018.10.038.

14. Miller BC, Sen DR, Al Abosy R, Bi K, Virkud YV, LaFleur MW, et al. Subsets of exhausted CD8(+) T cells differentially mediate tumor control and respond to checkpoint blockade. Nat Immunol 2019;20(3):326–36 doi 10.1038/s41590-019-0312-6.

15. Siddiqui I, Schaeuble K, Chennupati V, Fuertes Marraco SA, Calderon-Copete S, Pais Ferreira D, et al. Intratumoral Tcf1(+)PD-1(+)CD8(+) T Cells with Stem-like Properties Promote Tumor Control in Response to Vaccination and Checkpoint Blockade Immunotherapy. Immunity 2019;50(1):195–211 e10 doi 10.1016/j.immuni.2018.12.021.

16. Ficial M, Jegede OA, Sant’Angelo M, Hou Y, Flaifel A, Pignon JC, et al. Expression of T-Cell Exhaustion Molecules and Human Endogenous Retroviruses as Predictive Biomarkers for Response to Nivolumab in Metastatic Clear Cell Renal Cell Carcinoma. Clin Cancer Res 2021;27(5):1371–80 doi 10.1158/1078-0432.CCR-20-3084.

17. Detres Roman CR, Erwin MM, Rudloff MW, Revetta F, Murray KA, Favret NR, et al. Vaccination generates functional progenitor tumor-specific CD8 T cells and long-term tumor control. J Immunother Cancer 2024;12(10) doi 10.1136/jitc-2024-009129.

18. Beltra JC, Manne S, Abdel-Hakeem MS, Kurachi M, Giles JR, Chen Z, et al. Developmental Relationships of Four Exhausted CD8(+) T Cell Subsets Reveals Underlying Transcriptional and Epigenetic Landscape Control Mechanisms. Immunity 2020;52(5):825–41 e8 doi 10.1016/j.immuni.2020.04.014.

19. Cai HJ, Shi J, Yin LB, Zheng JF, Fu YJ, Jiang YJ, et al. Downregulation of TCF1 in HIV Infection Impairs T-cell Proliferative Capacity by Disrupting Mitochondrial Function. Front Microbiol 2022;13:880873 doi 10.3389/fmicb.2022.880873.

20. Ye Z, Gould TM, Zhang H, Jin J, Weyand CM, Goronzy JJ. The GSK3beta-beta-catenin-TCF1 pathway improves naive T cell activation in old adults by upregulating miR-181a. NPJ Aging Mech Dis 2021;7(1):4 doi 10.1038/s41514-021-00056-9.

21. Oliveira G, Stromhaug K, Klaeger S, Kula T, Frederick DT, Le PM, et al. Phenotype, specificity and avidity of antitumour CD8(+) T cells in melanoma. Nature 2021;596(7870):119–25 doi 10.1038/s41586-021-03704-y.

22. Schietinger A, Philip M, Krisnawan VE, Chiu EY, Delrow JJ, Basom RS, et al. Tumor-Specific T Cell Dysfunction Is a Dynamic Antigen-Driven Differentiation Program Initiated Early during Tumorigenesis. Immunity 2016;45(2):389–401 doi 10.1016/j.immuni.2016.07.011.

23. Ahuja D, Saenz-Robles MT, Pipas JM. SV40 large T antigen targets multiple cellular pathways to elicit cellular transformation. Oncogene 2005;24(52):7729–45 doi 10.1038/sj.onc.1209046.

24. Shiow LR, Rosen DB, Brdickova N, Xu Y, An J, Lanier LL, et al. CD69 acts downstream of interferon-alpha/beta to inhibit S1P1 and lymphocyte egress from lymphoid organs. Nature 2006;440(7083):540–4 doi 10.1038/nature04606.

25. D’Souza WN, Schluns KS, Masopust D, Lefrancois L. Essential role for IL-2 in the regulation of antiviral extralymphoid CD8 T cell responses. Journal of immunology 2002;168(11):5566–72 doi 10.4049/jimmunol.168.11.5566.

26. Schluns KS, Kieper WC, Jameson SC, Lefrancois L. Interleukin-7 mediates the homeostasis of naive and memory CD8 T cells in vivo. Nat Immunol 2000;1(5):426–32 doi 10.1038/80868.

27. Schluns KS, Williams K, Ma A, Zheng XX, Lefrancois L. Cutting edge: requirement for IL-15 in the generation of primary and memory antigen-specific CD8 T cells. Journal of immunology 2002;168(10):4827–31 doi 10.4049/jimmunol.168.10.4827.

28. Kawabe T, Yi J, Sprent J. Homeostasis of Naive and Memory T Lymphocytes. Cold Spring Harb Perspect Biol 2021;13(9) doi 10.1101/cshperspect.a037879.

29. Bamford RN, Grant AJ, Burton JD, Peters C, Kurys G, Goldman CK, et al. The interleukin (IL) 2 receptor beta chain is shared by IL-2 and a cytokine, provisionally designated IL-T, that stimulates T-cell proliferation and the induction of lymphokine-activated killer cells. Proc Natl Acad Sci U S A 1994;91(11):4940–4 doi 10.1073/pnas.91.11.4940.

30. Di Pilato M, Kfuri-Rubens R, Pruessmann JN, Ozga AJ, Messemaker M, Cadilha BL, et al. CXCR6 positions cytotoxic T cells to receive critical survival signals in the tumor microenvironment. Cell 2021;184(17):4512–30 e22 doi 10.1016/j.cell.2021.07.015.

31. Rathmell JC, Farkash EA, Gao W, Thompson CB. IL-7 enhances the survival and maintains the size of naive T cells. Journal of immunology 2001;167(12):6869–76 doi 10.4049/jimmunol.167.12.6869.

32. Kurtulus S, Tripathi P, Moreno-Fernandez ME, Sholl A, Katz JD, Grimes HL, et al. Bcl-2 allows effector and memory CD8+ T cells to tolerate higher expression of Bim. Journal of immunology 2011;186(10):5729–37 doi 10.4049/jimmunol.1100102.

33. Scott AC, Dundar F, Zumbo P, Chandran SS, Klebanoff CA, Shakiba M, et al. TOX is a critical regulator of tumour-specific T cell differentiation. Nature 2019;571(7764):270–4 doi 10.1038/s41586-019-1324-y.

34. Alfei F, Kanev K, Hofmann M, Wu M, Ghoneim HE, Roelli P, et al. TOX reinforces the phenotype and longevity of exhausted T cells in chronic viral infection. Nature 2019;571(7764):265–9 doi 10.1038/s41586-019-1326-9.

35. Khan O, Giles JR, McDonald S, Manne S, Ngiow SF, Patel KP, et al. TOX transcriptionally and epigenetically programs CD8(+) T cell exhaustion. Nature 2019 doi 10.1038/s41586-019-1325-x.

36. Yao C, Sun HW, Lacey NE, Ji Y, Moseman EA, Shih HY, et al. Single-cell RNA-seq reveals TOX as a key regulator of CD8(+) T cell persistence in chronic infection. Nat Immunol 2019;20(7):890–901 doi 10.1038/s41590-019-0403-4.

37. Akinduro O, Weber TS, Ang H, Haltalli MLR, Ruivo N, Duarte D, et al. Proliferation dynamics of acute myeloid leukaemia and haematopoietic progenitors competing for bone marrow space. Nature communications 2018;9(1):519 doi 10.1038/s41467-017-02376-5.

38. Choo DK, Murali-Krishna K, Anita R, Ahmed R. Homeostatic turnover of virus-specific memory CD8 T cells occurs stochastically and is independent of CD4 T cell help. Journal of immunology 2010;185(6):3436–44 doi 10.4049/jimmunol.1001421.

39. Miller I, Min M, Yang C, Tian C, Gookin S, Carter D, et al. Ki67 is a Graded Rather than a Binary Marker of Proliferation versus Quiescence. Cell Rep 2018;24(5):1105–12 e5 doi 10.1016/j.celrep.2018.06.110.

40. Roetman JJ, Erwin MM, Rudloff MW, Favret NR, Detres Roman CR, Apostolova MKI, et al. Tumor-Reactive CD8+ T Cells Enter a TCF1+PD-1-Dysfunctional State. Cancer Immunol Res 2023;11(12):1630–41 doi 10.1158/2326-6066.CIR-22-0939.

41. Escobar G, Tooley K, Oliveras JP, Huang L, Cheng H, Bookstaver ML, et al. Tumor immunogenicity dictates reliance on TCF1 in CD8(+) T cells for response to immunotherapy. Cancer Cell 2023;41(9):1662–79 e7 doi 10.1016/j.ccell.2023.08.001.

42. Peng Y, Fu S, Zhao Q. 2022 update on the scientific premise and clinical trials for IL-15 agonists as cancer immunotherapy. J Leukoc Biol 2022;112(4):823–34 doi 10.1002/JLB.5MR0422-506R.

43. Feng B, Bai Z, Zhou X, Zhao Y, Xie YQ, Huang X, et al. The type 2 cytokine Fc-IL-4 revitalizes exhausted CD8(+) T cells against cancer. Nature 2024;634(8034):712–20 doi 10.1038/s41586-024-07962-4.

44. Chen X, Hartman A, Guo S. Choosing Cell Fate Through a Dynamic Cell Cycle. Curr Stem Cell Rep 2015;1(3):129–38 doi 10.1007/s40778-015-0018-0.

45. Takahashi K, Yamanaka S. A decade of transcription factor-mediated reprogramming to pluripotency. Nat Rev Mol Cell Biol 2016;17(3):183–93 doi 10.1038/nrm.2016.8.

46. Staveley-O’Carroll K, Schell TD, Jimenez M, Mylin LM, Tevethia MJ, Schoenberger SP, et al. In vivo ligation of CD40 enhances priming against the endogenous tumor antigen and promotes CD8+ T cell effector function in SV40 T antigen transgenic mice. Journal of immunology 2003;171(2):697–707.

47. Stahl S, Sacher T, Bechtold A, Protzer U, Ganss R, Hammerling GJ, et al. Tumor agonist peptides break tolerance and elicit effective CTL responses in an inducible mouse model of hepatocellular carcinoma. Immunol Lett 2009;123(1):31–7 doi 10.1016/j.imlet.2009.01.011.

48. Brockstedt DG, Giedlin MA, Leong ML, Bahjat KS, Gao Y, Luckett W, et al. Listeria-based cancer vaccines that segregate immunogenicity from toxicity. Proc Natl Acad Sci U S A 2004;101(38):13832–7 doi 10.1073/pnas.0406035101.

49. Munoz-Ruiz M, Pujol-Autonell I, Rhys H, Long HM, Greco M, Peakman M, et al. Tracking immunodynamics by identification of S-G(2)/M-phase T cells in human peripheral blood. J Autoimmun 2020;112:102466 doi 10.1016/j.jaut.2020.102466.

50. Liboska R, Ligasova A, Strunin D, Rosenberg I, Koberna K. Most anti-BrdU antibodies react with 2’-deoxy-5-ethynyluridine -- the method for the effective suppression of this cross-reactivity. PLoS One 2012;7(12):e51679 doi 10.1371/journal.pone.0051679.

51. Espinosa-Carrasco G, Chiu E, Scrivo A, Zumbo P, Dave A, Betel D, et al. Intratumoral immune triads are required for immunotherapy-mediated elimination of solid tumors. Cancer Cell 2024 doi 10.1016/j.ccell.2024.05.025.

